# High-throughput colocalization pipeline quantifies efficacy of mitochondrial targeting signals across different protein types

**DOI:** 10.1101/2023.04.03.535288

**Authors:** Sierra K. Lear, Jose A. Nunez, Seth L. Shipman

## Abstract

Efficient metabolic engineering and the development of mitochondrial therapeutics often rely upon the specific and strong import of foreign proteins into mitochondria. Fusing a protein to a mitochondria-bound signal peptide is a common method to localize proteins to mitochondria, but this strategy is not universally effective with particular proteins empirically failing to localize. To help overcome this barrier, this work develops a generalizable and open-source framework to design proteins for mitochondrial import and quantify their specific localization. By using a Python-based pipeline to quantitatively assess the colocalization of different proteins previously used for precise genome editing in a high-throughput manner, we reveal signal peptide-protein combinations that localize well in mitochondria and, more broadly, general trends about the overall reliability of commonly used mitochondrial targeting signals.

## INTRODUCTION

Synthetic biologists increasingly leverage natural mitochondrial protein import pathways for compartmentalized metabolic engineering^1,2^ and the development of molecular therapeutics^3,4^. For metabolic engineering, sequestering enzymes within yeast mitochondria has resulted in a ∼300-fold increase in production of high-value biosynthetic compounds^5,6^. In human health, emerging mitochondrial therapeutics address a major unmet need since mutations to mitochondrial DNA (mtDNA) are at the root of numerous incurable diseases that affect over 1 in 5000 individuals^7,8^. In search of cures, researchers are exploring both allotopic expression, where corrected mitochondrial genes are expressed from the nuclear genome and sent to the mitochondria^9^, and gene editing, where mutated mtDNA is either depleted by nucleases or corrected by base editors^10-18^.

All of these approaches require efficient targeting of proteins of interest (POIs) to the mitochondria. The most common strategy to achieve such localization is by fusing a mitochondrial targeting sequence (MTS), typically a short and positively charged signal peptide, to the N-terminus of the POI. MTSs are recognized by translocases on the outer and inner mitochondrial membrane (TOM/TIM23 complex) that unfold the POI, import it through both mitochondrial membranes, and release the protein into the mitochondrial matrix following cleavage of the N-terminus MTS from the POI^19,20^. Hundreds of putative MTSs have been identified from natural proteins using computational tools^21–24^.

However, attachment of an individual MTS to a given POI does not always guarantee efficient import into mitochondria^25,3^. In fact, allotopic expression and gene editing approaches in mammalian mitochondria have been hindered by low or non-specific POI localization. For instance, only a small sub-selection of protein subunits typically encoded in mtDNA are able to be allotopically expressed in mammalian cells^26,27^. Moreover, even when proteins are imported to mammalian mitochondria, they can also accumulate in other organelles^28^. For mitochondrial gene editing, such imprecise localization poses danger; for example, the mitochondrial base editor DdCBE^16^ has substantial off-target editing in nuclear DNA^29^, highlighting the need for more specific import of genome editing-related proteins to mitochondria.

Previous studies in yeast suggest that length, hydrophobicity, charge, and folding of the MTS or POI can all affect the efficiency of mitochondrial import^30–32,19^. However, research in mammalian cells is much sparser and at present a given MTS-POI combination cannot be assumed to result in reliable mitochondrial import. Instead, researchers often empirically test multiple MTSs before finding one that results in their specific POI localizing in mitochondria^33,15^.

To address a relative lack of broad experimental data and help establish a quantitative assessment of mitochondrial localization in mammalian cells, we developed a quantitative and high-throughput imaging-based pipeline to measure POI import into mitochondria. Using this platform, we screened combinations of three commonly used N-terminus MTSs and POIs from five protein families relevant to mitochondrial gene editing to reveal the most reliable MTS-POI combinations.

## RESULTS

### High-throughput Localization Workflow

To investigate the effect of MTS on mitochondrial import across different POIs, we generated 66 protein cassettes containing a combination of localization signal and POI, followed by a HA tag on the C-terminus (**Fig 1a**). Localization signals included three commonly used MTSs—COX8 (29 amino acid-long peptide derived from human cytochrome c oxidase subunit VIII)^34^, Su9 (69 amino acid-long peptide derived from *Neurospora crassa* ATPase subunit 9)^35^, and ATG4D (42 amino acid-long peptide derived from a Atg4 cysteine protease)^36^—that were previously used to probe the import of CRISPR nucleases into mammalian mitochondria^15^. The use of no localization signal or a nuclear localization signal (NLS) served as negative controls. Nineteen POIs were chosen across five different protein classes that have been used as components of precise gene editing technologies: Class I CRISPR systems (Cas3/CASCADE)^37^, Class II CRISPR/Cas nucleases, RecTs, single-stranded binding proteins (SSBs), and retron reverse transcriptases (RTs). Both Class I and II CRISPR systems can cut DNA at programmable sites to induce editing using double-stranded break (DSB) repair pathways^38^. In contrast, RecTs and SSBs have been used to integrate donor DNA into bacterial and yeast genomes through recombineering^39–42^. Finally, retron RTs allow *in vivo* production of DNA donor editing templates in both prokaryotes and eukaryotes that mediate precise editing using either DSB repair pathways or recombineering^43–46^. As a positive control, we used the construct mito-APEX2, a protein that contains an MTS derived from the mitochondrially imported COX4 fused to APEX2, which has been shown to localize to the mammalian mitochondrial matrix using immunocytochemistry and proteomic mapping that found mito-APEX2 in close proximity to mitochondrial matrix proteins^47^.

**Figure 1.**
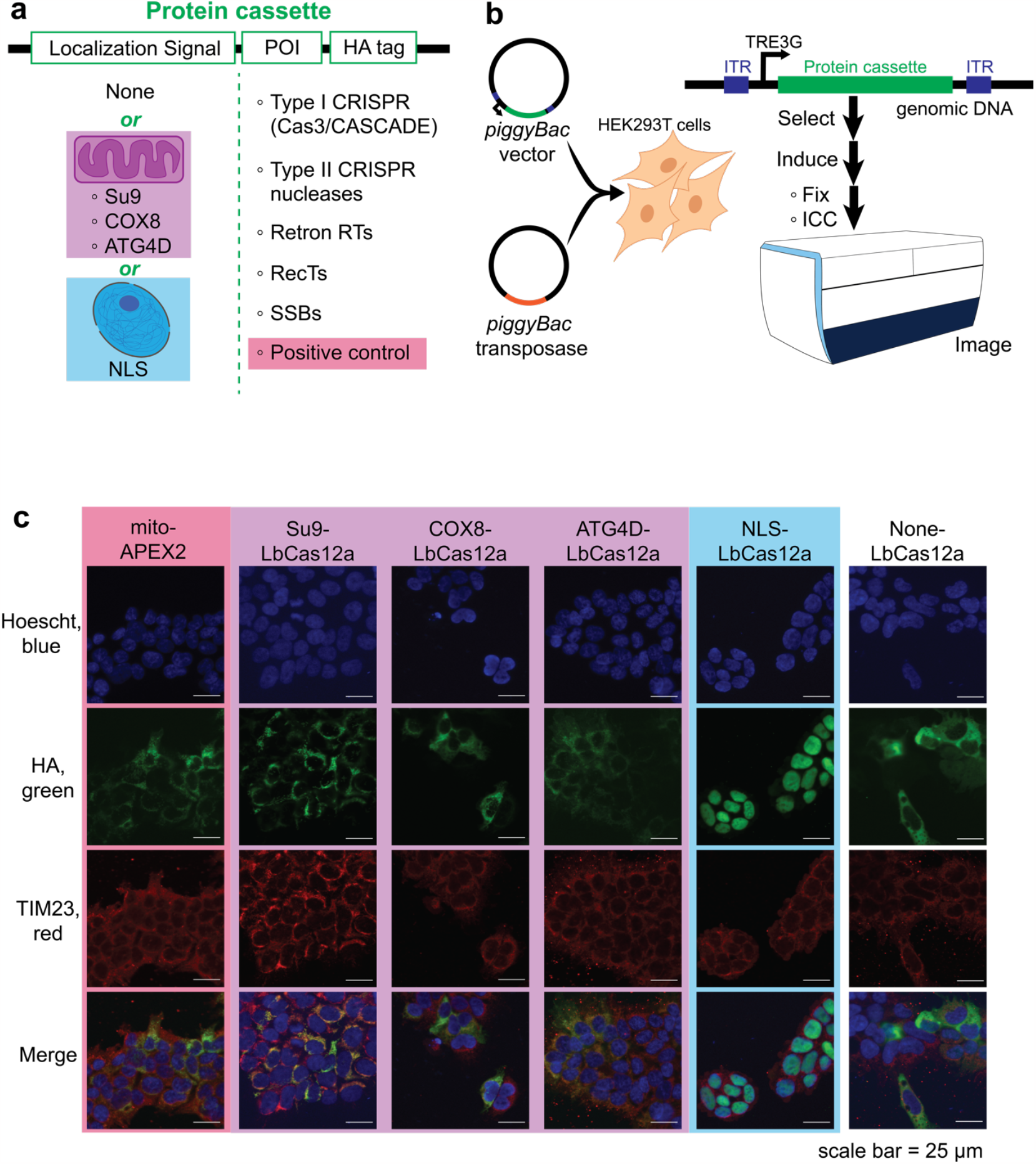
High-throughput localization workflow. **(a)** 66 genetic cassettes containing a combination of localization signal and POI, followed by a C-terminus HA tag, were synthesized. Localization signals include three N-terminus MTSs (Su9, COX8, and ATG4D), a nuclear localization signal (NLS), or no localization tag (None). POIs were chosen from five different protein families—Class I CRISPR/Cas proteins, Class II CRISPR nucleases, retron RTs, RecTs, and SSBs. A positive control of mito-APEX2 was also included. **(b)** Each cassette was randomly integrated into the genome of HEK293T cells using a piggyBac transposase system. Following co-transfection of the cassette in a piggyBac vector and a plasmid constitutively expressing piggyBac transposase, cells were selected for at least one week using the antibiotic puromycin. To image each cell line, expression of a given cassette was induced for 24 hours using doxycycline before being fixed and stained prior to imaging using a high-throughput confocal microscope (ImageXpress Micro Confocal High-Content Imaging System). **(c)** Engineered cell lines were stained with Hoescht (blue), an antibody against HA (green), and an antibody against the mitochondrial marker TIM23 (red). Shown are representative images from the positive control (red background) and LbCas12a fused to one of three MTSs (purple), NLS (blue), or no localization tag (white).

To engineer mammalian cell lines expressing a given protein cassette, each construct was cloned into a PiggyBac vector under the control of a doxycycline-inducible promoter adjacent to a constitutive puromycin resistance gene. These cassettes were randomly integrated into the genome of HEK293T cells using the PiggyBac transposase system and selected with puromycin (**Fig 1b**). Biological replicates of a given construct were defined as either individual clones derived from a single bulk transposase integration or multiple parallel transposase integrations. We screened the localization of each cassette by seeding cells into 96-well plates, expressed each protein for 24 hours under an inducible promoter, performed immunocytochemistry, and imaged each well using a high-throughput confocal microscope (ImageXpress Micro Confocal High-Content Imaging System). Specifically, each cell line was imaged for nuclei using Hoescht, POI using an antibody against HA, and mitochondria using an antibody against the mitochondrial marker TIM23 (**Fig 1c**). Using this high-throughput method, we found that localization of some cassettes varied qualitatively depending on localization signal. For instance, while the mito-APEX2 and Su9-LbCas12a showed punctate expression that colocalizes with mitochondria, NLS-LbCas12a showed clear nuclear colocalization and LbCas12a with no localization signal showed a diffuse, cytoplasmic phenotype. While these particular lines illustrate the expected localization based on signal, the localization of many other cassettes, such as of ATG4D-LbCas12a, were less predictable or more ambiguous. Thus, we next developed an analytical pipeline to quantify localization within mitochondria or nuclei.

### Automated, quantitative, and open-source analysis pipeline

To better compare localization differences exhibited by MTS-POI combinations, we developed an unbiased Python-based analysis pipeline to quantify the mitochondrial and nuclear import between our dozens of cassettes. Crucially, we found that expression and localization were variable between individual cells of a given condition so our analysis pipeline is built to quantify colocalization at the level of single cells. Images corresponding to the nuclei and mitochondria for each biological replicate cell line from each condition were fed into a Cellpose-based machine learning model^48,49^ to label individual cells (**Fig 2a,b**). Next, cells were filtered using Otsu thresholding to remove any cells with no detectable protein expression (**Fig 2c**). Specifically, Otsu thresholding was applied on each image to determine the pixel intensity threshold separating POI signal from background fluorescence. This analysis also revealed that some images had such low fluorescence that signal was effectively indistinguishable from noise. To ensure these specific images did not bias the final colocalization scores, any images in which Otsu thresholding did not separate signal from noise in each cell, as defined by the majority of filtered cells in an image failing to show a non-Gaussian intensity distribution typical of true fluorescent signal, were discarded (**Fig S1**). In some cases, only a few images for a cell line were eliminated, although—in cases where a clonal or transfected line suffered from minimal cassette expression—the entire biological replicate was removed from analysis.

**Figure 2.**
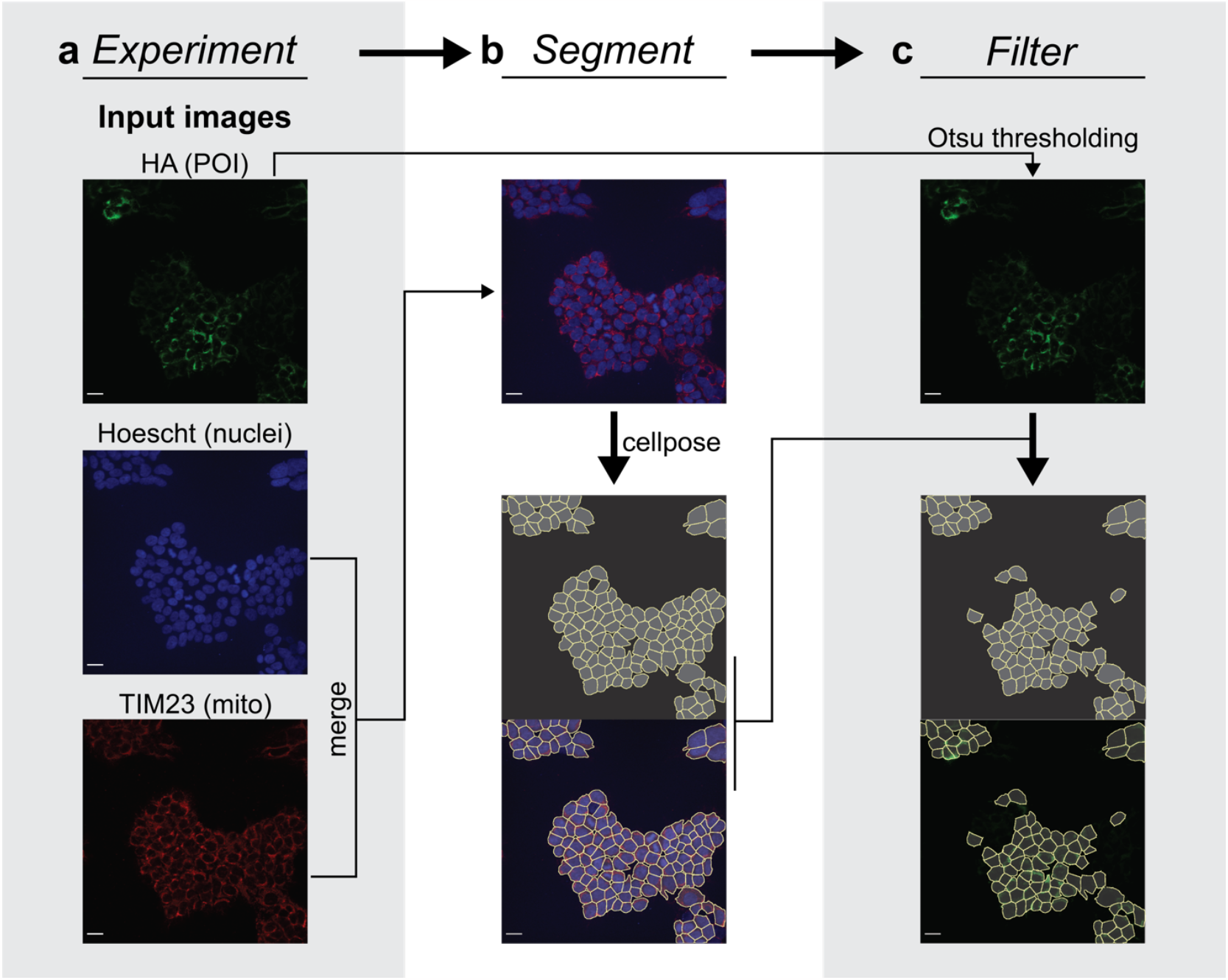
Machine learning algorithm enables automated labeling of cells containing cassette protein. Scale bar = 25 μm. **(a)** Unprocessed images from each fluorescent channel are fed into a Python-based analysis pipeline to segment individual cells. Shown are representative images from the cell line Su9-LbCas12a (top, HA; middle; Hoescht, bottom; TIM23). **(b)** The Hoescht and TIM23 channels are merged (top) prior to being fed into a custom, retrained neural network (arrow), resulting in automated labeling of each cell found within the image. Images below arrow; top image shows Cellpose-generated mask of segmented cells, where each yellow line indicates a cell boundary. Bottom image shows mask overlaid atop merged Hoescht/TIM23 channels. **(c)** Cellpose-segmented cells are filtered to keep only cells which express the cassette protein. Otsu thresholding is applied to the HA channel (top) to determine a threshold separating true fluorescent signal from background noise. Segmented cells containing at least 50 pixels with signal are kept to perform further colocalization measurements. Images below arrow; top image shows segmented cells following filtering. Bottom image shows filtered mask overlaid atop HA channel.

After selecting a population of filtered cells to further analyze for a given protein cassette, colocalization between the cassette protein and either mitochondria (**Fig 3a,b**) or nuclei (**Fig 3d,e**) was measured on a per cell basis using Pearson’s correlation coefficient (PCC)^50^. Using this method, PCC scores vary between -1 (anti-correlated) to +1 (highly correlated). High colocalization scores indicate that a protein is collocated with a given organelle while low colocalization scores suggest little to no specific colocalization between a POI and a given organelle occurred.

**Figure 3.**
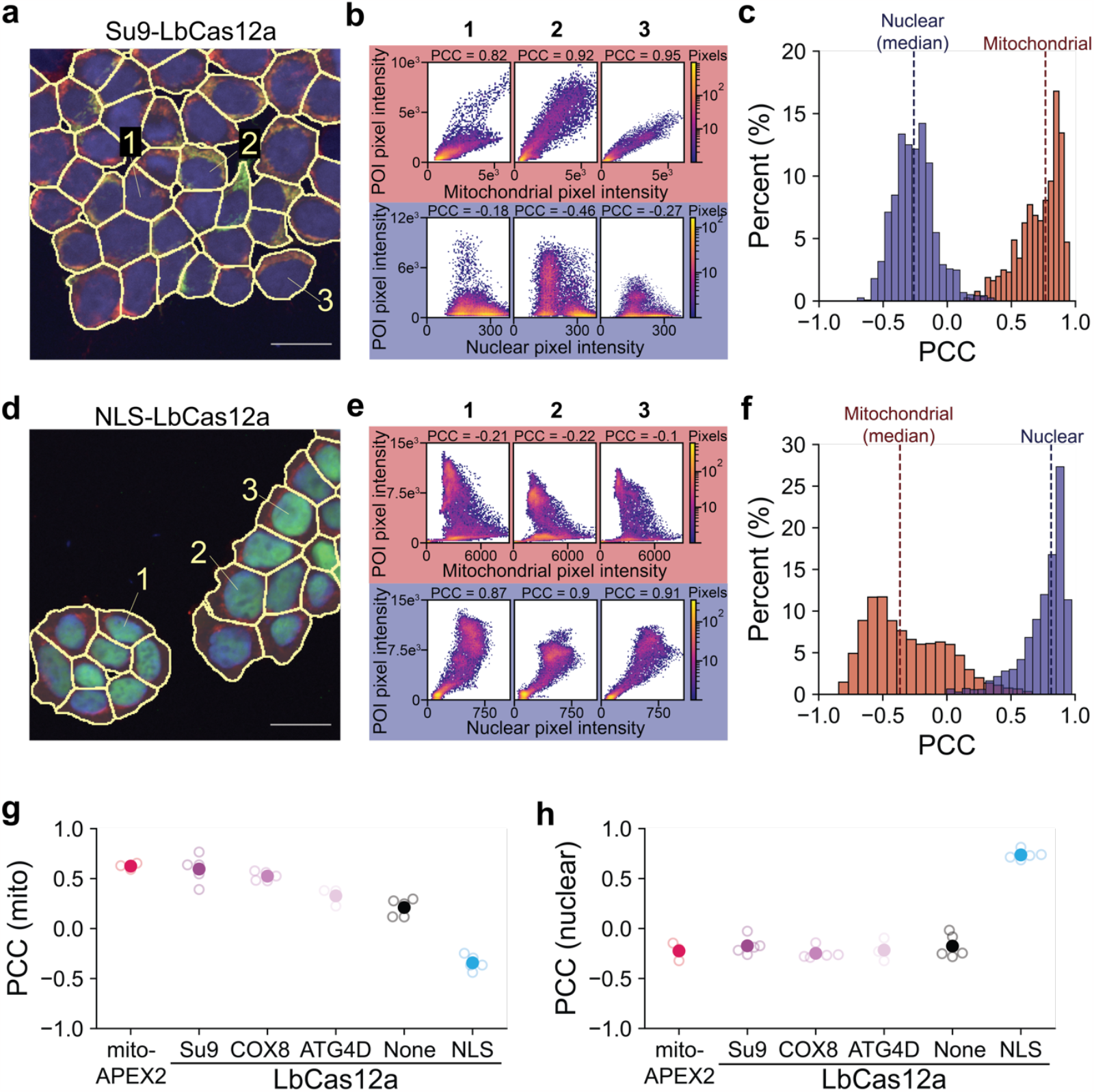
Computational workflow quantifies the colocalization of cassette proteins with mitochondria or nuclei. **(a)** Representative image of a cell line (Su9-LbCas12a) with clear mitochondrial expression of its cassette protein, with mask in yellow overlaid on top. Numbers refer to three representative cells for which data is shown in (b). Scale bar = 25 μm. **(b)** Heatmaps depicting the relationship between Su9-LbCas12a pixel intensity and organellar pixel intensity (mitochondria on top in red; nuclei on bottom in blue) for each pixel within a representative cell from (a) (left; cell #1, middle; cell #2, right; cell #3). Color depicts the number of pixels. The strength of the linear relationship between pixel intensities, or colocalization, within each cell is calculated using PCC, and the result depicted on top its respective heatmap. **(c)** Histogram depicting the all the colocalization scores for all the cells for one clonal line expressing Su9-LbCas12a. Mitochondrial PCC scores are shown in red, while nuclear PCC scores are shown in blue. Dotted lines depict the median colocalization score for mitochondria (red) and nuclei (blue). **(d)** Representative image of a cell line (NLS-LbCas12a) with clear nuclear expression of its cassette protein, with mask in yellow overlaid on top. Numbers refer to three representative cells. Scale bar = 25 μm. **(e)** Heatmaps depicting the relationship between NKS-LbCas12a pixel intensity and organellar pixel intensity (mitochondria on top in red; nuclei on bottom in blue) for each pixel within a representative cell from (d) (left; cell #1, middle; cell #2, right; cell #3). Color depicts the number of pixels. The strength of the linear relationship between pixel intensities, or colocalization, within each cell is calculated using PCC, whose result is on top its respective heatmap. **(f)** Histogram depicting the all the colocalization scores for all the cells for one clonal line expressing NLS-LbCas12a. Mitochondrial PCC scores are shown in red, while nuclear PCC scores are shown in blue. Dotted lines depict the median colocalization score for mitochondria (red) and nuclei (blue). **(g)** Mitochondrial colocalization of positive control mito-APEX2 (red) and LbCas12a fused to different localization tags, as measured using the described experimental and analytic workflow. There is a significant effect of localization signal (one-way ANOVA, *P*<0.0001), where ATG4D (*P*=0.0021), no signal (*P*<0.0001), and NLS (*P*<0.0001) are all significantly different from mito-APEX2, but Su9 (*P*=0.9831) and COX8 (*P*=0.376) are not (Dunnett’s corrected). Open circles are biological replicates; closed circles are average of all biological replicates. **(h)** Nuclear colocalization of mito-APEX2 (red) and LbCas12a fused to different localization tags measured using the described experimental and analytical workflow. There is a significant effect of localization signal (one-way ANOVA, *P*<0.0001), where NLS (*P*<0.0001) is significantly different from mito-APEX2, but Su9 (*P*=0.8587), COX8 (*P*=0.9930), ATG4D (*P*=0.9998), and no signal (*P*=0.0.8825) are not (Dunnett’s corrected). Open circles are biological replicates; closed circles are average of all biological replicates. Additional statistical details in Supplementary Table 1.

For our analysis, we considered individual cells that survived quality filters from a single transfection or clone as technical replicates and summarized the overall colocalization score for a single biological replicate of each protein cassette by taking the median of all the individual cell colocalization scores for mitochondria (**Fig 3c**) or nuclei (**Fig 3f**). We replicated our experiments using at least three different transfections or five clonal lines as biological replicates.

We generally found low variability within our biological replicates, suggesting that protein import is a fairly reliable phenomenon. The positive control mito-APEX2 obtained an average score of 0.63 +/-0.03 (mean +/-std. dev), a high PCC value that strongly implies mito-APEX2 is imported into the mitochondrial matrix. Similar to our previous qualitative assessments (**Fig 1c**), we found that colocalization scores, even across a single POI, vary depending on localization signals. The colocalization scores of LbCas12a fused to the Su9 or COX8 MTSs were not statistically different from mito-APEX2, suggesting mitochondrial import had occurred. In comparison, when LbCas12a was instead fused to ATG4D, no localization signal, or NLS, colocalization scores dropped significantly, indicating less mitochondrial import occurred (**Fig 3g**). Moreover, when comparing nuclear colocalization scores (**Fig 3h**), all cell lines except NLS-LbCas12a showed a consistent, low nuclear colocalization score, suggesting little to no nuclear import. As expected, only the cell line fused to a NLS had a high colocalization score indicating high nuclear import. These findings suggest that our workflow is able to compare import efficiencies across different combinations of MTS and POI for multiple organelles.

### MTS selection strongly influences mitochondrial import

After validating the analytical pipeline, we used this workflow to quantify the mitochondrial and nuclear import of all 66 different protein cassettes (**Fig 4a; S2a**,**b**). Interestingly, mitochondrial colocalization scores did not cluster bimodally into high and low scores. Instead, scores were distributed continuously, suggesting that different cassettes have varying capabilities to drive POIs to the mitochondria.

**Figure 4.**
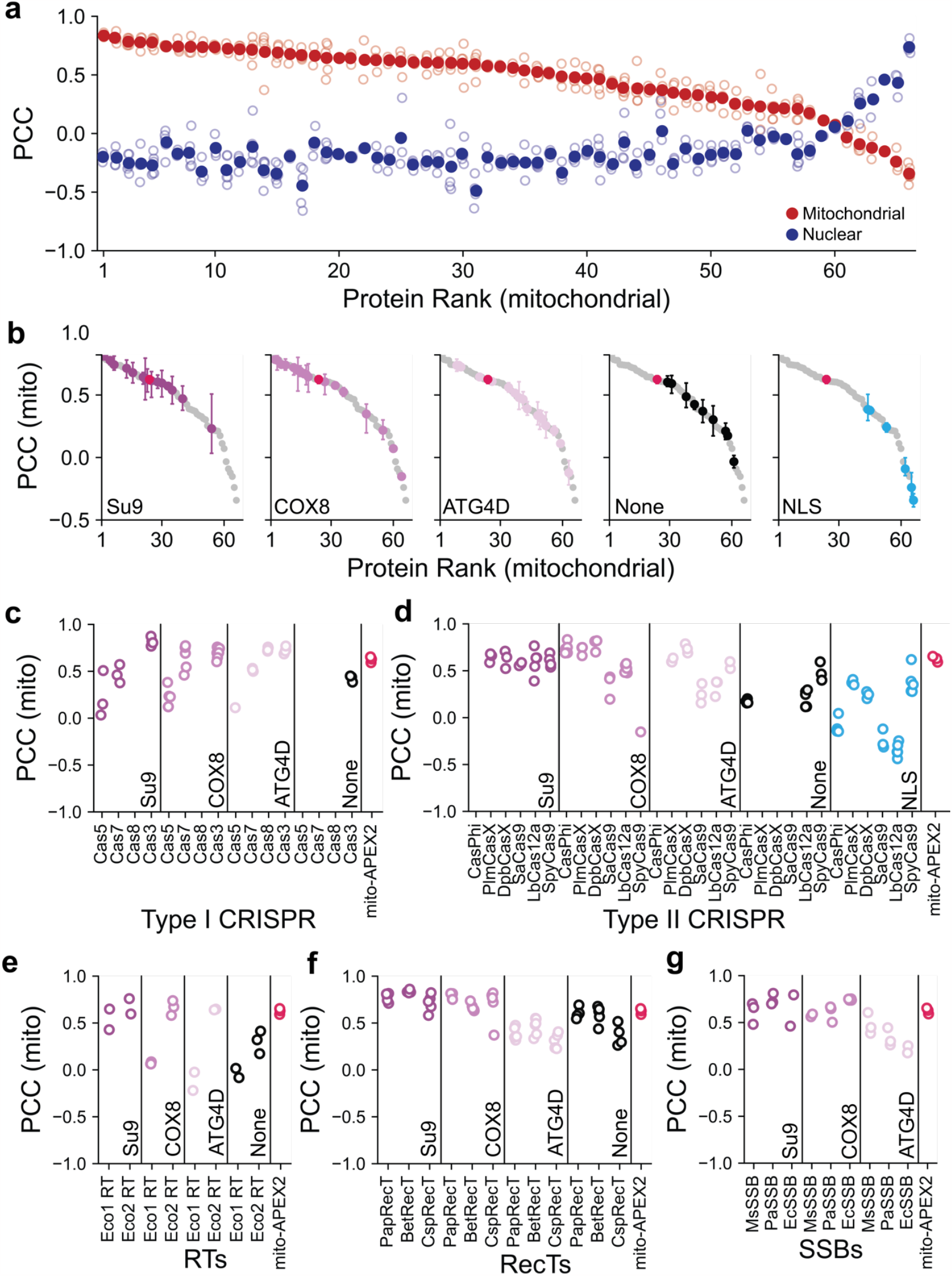
MTS selection strongly influences mitochondrial import. **(a)** Mitochondrial (red) and nuclear (blue) colocalization scores of 66 different protein cassettes. Proteins were ranked based on average mitochondrial colocalization score, from highest mitochondrial PCC to lowest. Open circles are biological replicates; closed circles are average of all biological replicates. **(b)** Clustering of cassettes based on specific localization signal. The entire continuum of all mitochondrial colocalization scores from (a) is shown in each subplot in gray, while the positive control mito-APEX2 is shown in each subplot in red. The clustering of cassettes with a specific localization signal is shown in another color on top (from left to right subplots: Su9, COX8, ATG4D, none, and NLS). There is a significant effect of localization signal (linear mixed effects model, *P*<0.0001), where ATG4D is significantly different from both Su9 (*P*<0.0001) and COX8 (*P*<0.0001) but not None (0.092559) during follow-up (Kenward-Roger corrected). Meanwhile, there is no significant difference between Su9 and COX8 (*P*=0.88139). Closed circles indicate average of all biological replicates; error bars indicate standard deviation. **(c)-(g)** Mitochondrial colocalization scores broken down by localization signal within a given protein family (**c**; Class I CRISPR Cas3/CASCADE, **d**; Class II CRISPR nucleases, **e**; retron RTs, **f**; RecTs, **g**; SSBs). Specific proteins are listed in order of amino acid length, from shortest on the left to longest on the right. As a reference for mitochondrial PCC scores suggestive of mitochondrial import, scores for positive control mito-APEX2 are shown to the right of each figure (red). Open circles are individual biological replicates.

Previous studies have found that both the specific MTS and hydrophobicity of individual POIs can influence the probability that a MTS-POI will localize in the mitochondria ^31,19,33^. To determine if localization signal influences mitochondrial import across the four protein families tested, we investigated how the mitochondrial import rank of cassettes clustered based on localization signal (**Fig 4b**). We fit a linear mixed effects model with a fixed effect of localization signal and a random intercept within each protein family, to specifically test the statistical significance of each localization signal while accounting for the variability inherent within each protein of interest. As expected, we found that cassettes clustered based on specific localization signals; while fusion to an MTS resulted in higher rankings across the board, having a NLS led to clear clustering at the bottom of the rankings. However, there was a clear ranking priority to how well each MTS performed compared to each other, with Su9 and COX8 driving high mitochondrial import, while ATG4D performed significantly worse.

Given ATG4D’s uneven performance across different POIs, we next further analyzed how colocalization scores varied based on localization scores within different protein families (**Fig 4c-g**). Interestingly, all three MTSs were able to drive high mitochondrial import of Class I CRISPR-related proteins (**Fig 4c**), whereas both COX8 and ATG4D appeared less capable than Su9 of importing the larger Class II CRISPR nucleases (**Fig 4d**). Of the two retron RTs tested, only Su9 led to mitochondrial localization of retron-Eco1 RT, whereas COX8 and ATG4D were able to localize retron Eco2 RT (**Fig 4e**). Finally, while both COX8 and Su9 were able to import all RecTs and SSBs tested into the mitochondria (**Fig 4f,g**), ATG4D instead appeared to misdirect these proteins to a different location, based on an unusual punctate pattern which did not colocalize with mitochondria (**Fig S2c)**.

## DISCUSSION

Importing non-canonical proteins into mammalian mitochondria is a critical step to overcome particular challenges in metabolic engineering and for developing therapeutics to treat genetic diseases of the mtDNA. Here, we design a high-throughput imaging-based workflow to quickly screen the subcellular localization of a tagged protein. This method enabled us to determine the best MTS-POI combination across three commonly used MTSs and five different protein classes that are components of gene editing technologies. The results from these screens should help drive more robust mitochondrial gene editing by enabling researchers to test mitochondrial gene editing using other nucleases or proteins beyond SpyCas9, which has been reported to cause mitochondrial dysfunction when imported to mitochondria^15^, or testing MTSs that yield more specific mitochondria import to prevent off-target effects in nuclear DNA^29^.

This work also reveals broader trends about which MTS to use across multiple protein types. By testing three common MTSs on a diverse set of POIs, we find that Su9 and COX8 consistently performed the best while ATG4D performed the most unevenly and misdirected several POIs to an alternate location. In addition, unlike COX8 and ATG4D, Su9 was able to import specific proteins, such as large Class II nucleases and Eco1 RT, into the mitochondria.

As a tool for the field, our Python-based workflow implemented in an annotated Jupyter notebook can be reused for future experiments, including more generally to other screens using different proteins, organelles of interest, or cell lines. Although we used a high-throughput confocal microscope in our own workflow, other confocal microscopes could easily be used, depending on the necessary throughput or number of samples. Computationally, the analytic pipeline uses Python, an open-source programming language, and relies on pixel-based colocalization analyses and quality check steps to eliminate cells not expressing a protein of interest that can be universally applied regardless of cell line or differing expression levels or phenotypes^50^. The only experiment-specific alteration to the pipeline would be to apply a different neural network to segment individual cells, depending on the cell line and seeding density used. However, Cellpose already offers a collection of ready-to-use neural networks or—if further refinement is necessary—an intuitive GUI that enables users to retrain and create their own neural network in fewer than 30 minutes^49^. Therefore, others should be able to easily apply this analytical framework to quantify colocalization within their own fluorescent images.

## Methods

### Constructs and strains

Protein cassettes were constructed by amplifying localization signals and POI nucleotide sequences using PCR from synthesized gBlocks (IDT) or existing plasmids. Complete protein cassettes were cloned into a PiggyBac integrating plasmid for doxycycline-inducible human protein expression (TetOn-3G promoter) using Gibson assembly. Alternatively, some cassettes were synthesized into the same custom PiggyBac integrating plasmid by Twist Bioscience (see Supplementary Table 2).

Stable mammalian cells for imaging were generated using the standard Lipofectamine 3000 transfection protocol (Invitrogen) and a PiggyBac transposase system. T12.5 flasks with 50-70% confluent HEK293T cells were transfected using 1.6 μg POI cassette expression plasmid and 0.8 μg PiggyBac transposase plasmid (pCMV-hyPBase). Stable cell lines were selected using puromycin for at least one week.

Clonal lines were generated by growing individual cells into separate cell populations. Specifically, stable cell lines were serially diluted to a final concentration of 2.5 cells per mL media then seeded into a 96-well plate using 100 μL/well. Wells that received a single cell had media refreshed weekly until a clonal line proliferated to ∼40% confluency, at which point a clonal line was passaged to a larger flask for further experiments.

### Immunocytochemistry

96-well glass bottom plates with #1 cover glass (Cellvis, catalog # P96-1-N) were coated with a mixture of 50% poly-D-lysine (ThermoFisher Scientific, catalog #A3890401) and DPBS (ThermoFisher Scientific, catalog #14040133) for 30 minutes at room temperature. Wells were washed three times with distilled water and left out to dry for at least 2 hours prior to seeding.

Cells were seeded at a density of 10,000 cells per well. The following day, doxycycline was added at a final concentration of 1 μg/mL to induce expression of the protein cassette. At 24 hours post-induction, cell nuclei were stained using a final concentration of 10 μM Hoescht for at least 5 minutes prior to fixation.

For fixation, media was aspirated from each well and replaced with a solution of 4% paraformaldehyde (PFA) created fresh by fixing a 1 mL 16% (w/v) PFA ampule (ThermoFisher Scientific, catalog #28906) with 3 mL PBS. Cells were fixed for 30 minutes at room temperature prior to three 5-minute washes with PBS. Following fixation, cells were permeabilized and blocked for an hour at room temperature using blocking buffer made fresh with the following ingredients: PBS containing 10% donkey serum (Sigma-Aldrich, catalog #D9663), 10% Triton X-100 (Sigma-Aldrich, catalog #X100), and 100 mg BSA (Sigma-Aldrich, catalog #A9418) per 10 mL solution. Next, cells were incubated overnight at 4°C in blocking buffer with the antibodies anti-HA tag conjugated to DyLight 550 (ThermoFisher Scientific, catalog #26183-D550) and anti-TIM23 (Abcam, catalog #ab230253) each added at a 1:100 dilution. After performing three more 5-minute washes, cells were incubated with a secondary antibody goat anti-rabbit conjugated to DyLight 650 (ThermoFisher Scientific, catalog #84546) at a 1:500 dilution in blocking buffer for 3 hours. Following secondary antibody incubation, three more 5-minute washes were performed prior to the addition of 30 uL antifade mountant (ThermoFisher Scientific, catalog #S36967) per well.

Plates were wrapped in aluminum foil to avoid light and either stored temporarily at 4°C or at -20°C for longer-term storage prior to imaging.

### Imaging

Stained cells were imaged using an ImageXpress Micro Confocal High-Content Imaging System (Molecular Devices) using a 40X water immersion objective by taking a 7-layer Z-stack, with each layer spaced 0.3 μm apart, at four different sites per well.

### Colocalization Image Analysis Pipeline

A colocalization image analysis pipeline was made using jupyter-notebook in Python 3^51,52^, and uses the following packages: numpy, pandas, scipy, skimage, tdqm, and tifffile. Additionally, the pipeline requires the Cellpose code library^48,49^ along with these additional packages: numba, opencv, and pytorch. Using the Cellpose GUI also requires PyQt and pyqtgraph^53^.

TIFF files consisting of merged nuclear and mitochondrial channels were created using a custom function and fed into a neural network retrained according to the instructions for the Cellpose GUI^49^. Briefly, the “CP” model from the Cellpose model zoo was initially used to segment all images. Afterwards, about five images with poor initial segmentation were chosen for manual annotation. The CP model was then retrained using the corrected labels and the new model was re-run on all images.

To remove segmented cells that did not contain expression of the cassette protein, Otsu thresholding was performed on the HA channel to determine a pixel intensity threshold separating signal from background for each image. Only segmented cells containing at least 50 pixels of cassette protein signal, referred to as filtered cells, were kept for further analysis.

Two additional functions to ensure quality-check steps were also implemented. First, any image containing fewer than six filtered cells was automatically removed from further analysis. Second, since the overall expression of a cassette protein can vary between different cell lines, a function was written to ensure that the filtering step effectively distinguished between cells that did or did not express a cassette protein. Individual cells containing noise, rather than signal, exhibit a Gaussian distribution of protein cassette pixel intensities. In contrast, cells with signal tend to exhibit non-normal or skewed pixel intensity distributions. Thus, for every image, a “non-Gaussian” test was performed on each filtered cell by testing for normality. If over 60% of filtered cells failed the “non-Gaussian” test, then this result suggests that the majority of filtered cells within the image do not contain true expression of the cassette protein, thus that specific image would be removed from further analysis.

Afterwards, a custom function was built to calculate PCC between the HA channel pixel intensities and either the mitochondrial or nuclear channel pixel intensities for every filtered cell related to a given biological replicate. Due to the skew present in most PCC distribution, these results were summarized by taking the median of all the filtered cells for a given biological replicate.

### Statistics

ANOVA and post-hoc analyses were performed using GraphPad Prism v9.4.1 For linear mixed effects modeling, we used *R* v4.2.3 using the *lme4* and *lmerTest* packages (post-hoc analyses used the Kenward-Roger degrees of freedom correction method implemented in the package *pbkrtest*).

## ABBREVIATIONS

MTS: mitochondrial targeting sequence
POI: protein of interest
mtDNA: mitochondrial DNA
CRISPR: clustered regularly interspaced short palindromic repeats

## Acknowledgements

This work was supported by the National Institute of General Medical Sciences (DP2GM140917), the National Institute of Biomedical Imaging and Bioengineering (R21EB031393), and the UCSF Program for Breakthrough Biomedical Research (partially funded by the Sandler Foundation). S.K.L. was supported by an NSF Graduate Research Fellowship (2034836). S.L.S. is a Chan Zuckerberg Biohub – San Francisco Investigator. We thank David Darevsky for his advice on coding and statistics.

