## Supporting Information for "High-throughput colocalization pipeline quantifies efficacy of mitochondrial targeting signals across different protein types"

**Contents:**

**Supplemental Figures 1 and 2**
**Supplemental Tables 1 and 2**
**Supplemental References**

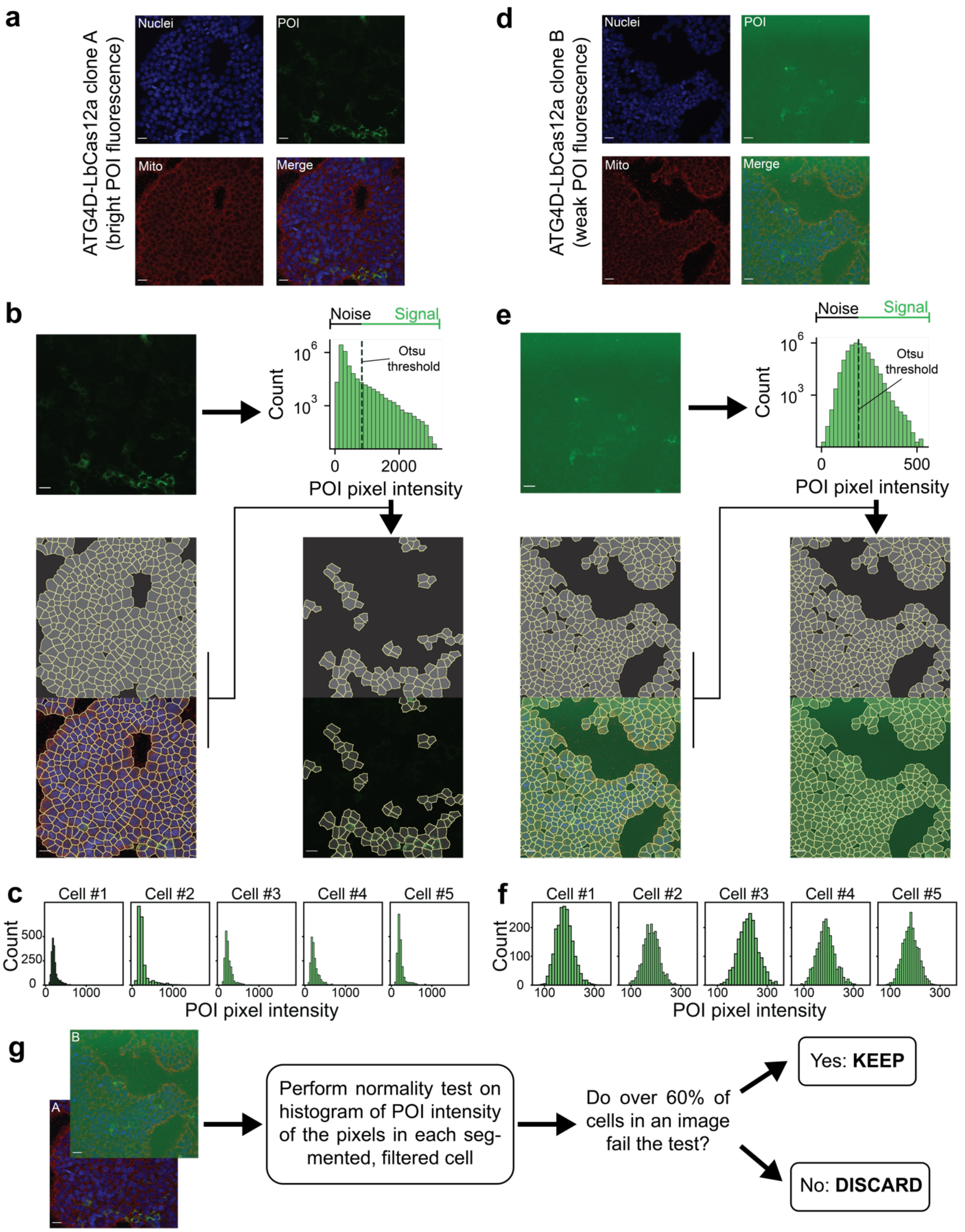

**Supplemental Figure 1.** Corresponds to Figure 2 in the main text. Normality test implemented in Python to remove cell lines with low signal-to-noise ratio of cassette protein, as part of a computational workflow to quantify the import of cassette proteins into mitochondria or nuclei. All scale bars = 25 µm. **(a)** Representative image of a clonal line of ATG4D-LbCas12a with strong expression of its cassette protein across three fluorescent channels (Hoescht, TIM23, and HA) and a merge of all channels. **(b)** Segmenting and filtering using Otsu threshold for the representative image in (a). Otsu thresholding is applied to determine pixel intensity threshold that separates "noise" from "signal" or true fluorescence. This threshold is used to remove all individual cell masks that do not have >50 pixels of "signal" to create a smaller sub-selection of filtered cells to analyze. **(c)** To check that the filtering algorithm appropriately removed cells without signal, the shape of the histogram of POI intensity pixels per filtered cell is checked for normality. Here, give representative cells from the filtered cells chosen in (b) are shown. Note all histograms show a skewed, non-normal shape that is indicative of true signal. **(d)** Representative image of clonal line of ATG4D-LbCas12a with weak expression of its cassette protein across three fluorescent channels (Hoescht, TIM23, and HA) and a merge of all channels. **(e)** The same segmenting and filtering process and described in (b) is shown for the representative image from (d). Note that the chosen Otsu threshold does not eliminate any cells from the mask. **(f)** The histograms of POI pixel intensity from five representative cells from the filtered cells from (e) are shown. Note all histograms show a normal distribution, indicative of noise rather than true signal. **(g)** Schematic showing the implementation of normality test to remove cell lines with low signal-to-noise ratio of cassette protein. A normality test on the histogram of POI intensity in each filtered cell is performed per image. If over 60% of the cells in an image are non-normal, indicative of true signal, the image is kept while. If over 60% of the cells are normal, then the image is instead discarded and not included in future analyses.

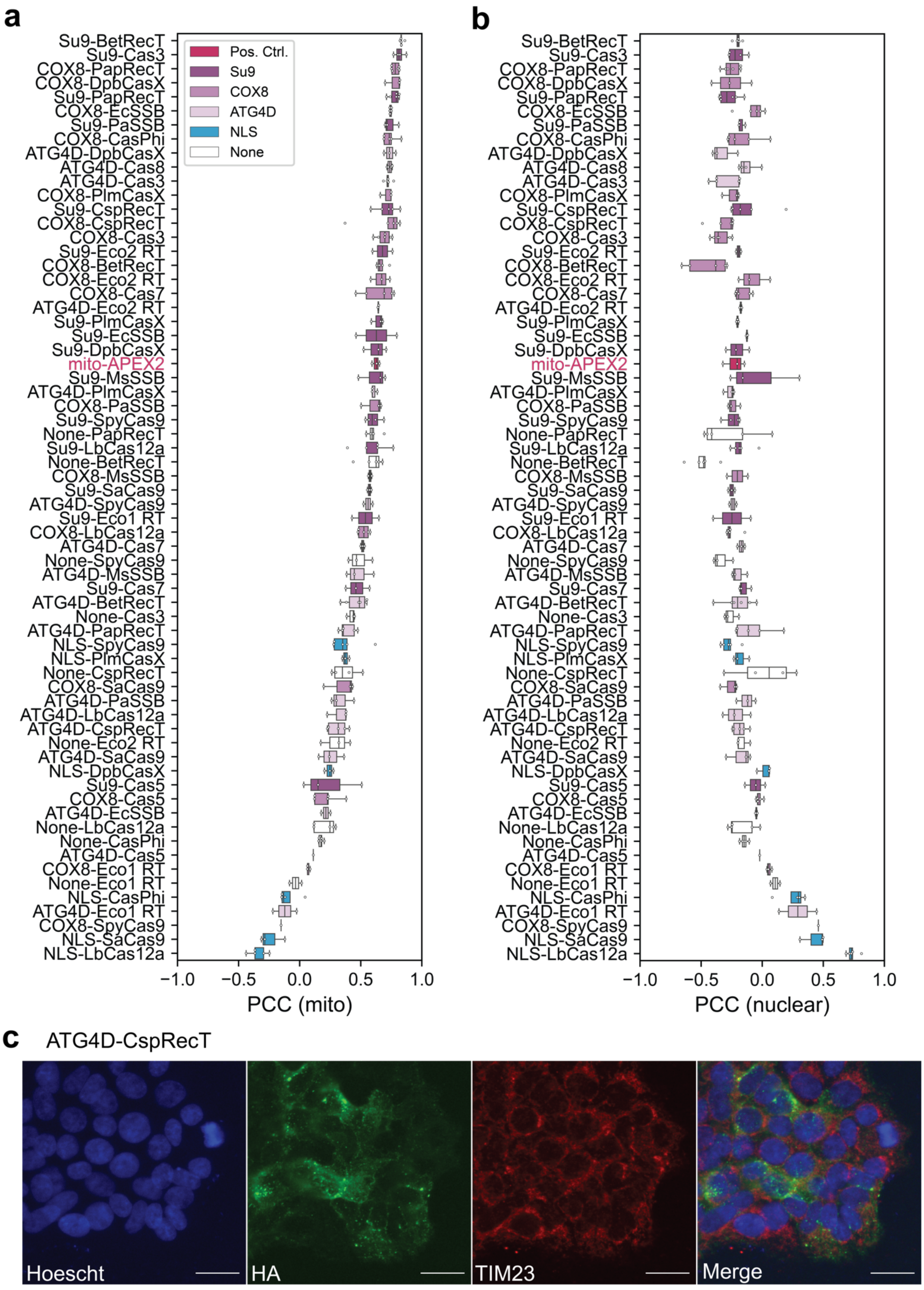
a
Pos. Ctrl.
Su9
COX8
ATG4D
NLS
None
Su9-BetRecT
Su9-Cas3
COX8-PapRecT
COX8-DpbCasX
Su9-PapRecT
COX8-EcSSB
Su9-PaSSB
COX8-CasPhi
ATG4D-DpbCasX
ATG4D-Cas8
ATG4D-Cas3
COX8-PlmCasX
Su9-CspRecT
COX8-CspRecT
COX8-Cas3
Su9-Eco2 RT
COX8-BetRecT
COX8-Eco2 RT
COX8-Cas7
ATG4D-Eco2 RT
Su9-PlmCasX
Su9-EcSSB
Su9-DpbCasX
mito-APEX2
Su9-MsSSB
ATG4D-PlmCasX
COX8-PaSSB
Su9-SpyCas9
None-PapRecT
Su9-LbCas12a
None-BetRecT
COX8-MsSSB
Su9-SaCas9
ATG4D-SpyCas9
Su9-Eco1 RT
COX8-LbCas12a
ATG4D-Cas7
None-SpyCas9
ATG4D-MsSSB
Su9-Cas7
ATG4D-BetRecT
None-Cas3
ATG4D-PapRecT
NLS-SpyCas9
NLS-PlmCasX
None-CspRecT
COX8-SaCas9
ATG4D-PaSSB
ATG4D-LbCas12a
ATG4D-CspRecT
None-Eco2 RT
ATG4D-SaCas9
NLS-DpbCasX
Su9-Cas5
COX8-Cas5
ATG4D-EcSSB
None-LbCas12a
None-CasPhi
ATG4D-Cas5
COX8-Eco1 RT
None-Eco1 RT
NLS-CasPhi
ATG4D-Eco1 RT
COX8-SpyCas9
NLS-SaCas9
NLS-LbCas12a
−1.0 −0.5 0.0 0.5 1.0
PCC (mito)
b
−1.0 −0.5 0.0 0.5 1.0
PCC (nuclear)
c
ATG4D-CspRecT
Hoescht
HA
TIM23
Merge

**Supplemental Figure 2.** Corresponds to Figure 4 in the main text. Summary of the import of protein cassettes using our quantitative, high-throughput pipeline. **(a)** Mitochondrial colocalization scores of 66 different protein cassettes. Proteins are arranged from top to bottom based on average mitochondrial colocalization score, from highest mitochondrial PCC to lowest. Open circles are technical replicates. **(b)** Nuclear colocalization scores of 66 different protein cassettes. Proteins are arranged from top to bottom based on average mitochondrial colocalization score, from highest mitochondrial PCC to lowest. Open circles are technical replicates. **(c)** Representative image of a clonal line of ATG4D-CspRecT across three fluorescent channels (Hoescht, HA, and TIM23) and a merge of all channels. Note the punctate expression of HA that does not align with mitochondria (TIM23).

### Supplementary Table 1, Statistics

| Figure 3 | | | | | |
|---|---|---|---|---|---|
| **Panel** | **Biological replicates** | **Comparison** | **Test** | ***P* value** | ***P* value summary** |
| **g** | 3-5 | **PCC (mito): effect of condition** | **one-way ANOVA** | <0.0001 | **** |
| | | Follow-up: mito-APEX2 vs. Su9-LbCas12a | Dunnett's multiple comparison test (corrected) | 0.9831 | ns |
| | | Follow-up: mito-APEX2 vs. COX8-LbCas12a | Dunnett's multiple comparison test (corrected) | 0.3760 | ns |
| | | Follow-up: mito-APEX2 vs. ATG4D-LbCas12a | Dunnett's multiple comparison test (corrected) | 0.0021 | ** |
| | | Follow-up: mito-APEX2 vs. None-LbCas12a | Dunnett's multiple comparison test (corrected) | <0.0001 | **** |
| | | Follow-up: mito-APEX2 vs. NLS-LbCas12a | Dunnett's multiple comparison test (corrected) | <0.0001 | **** |
| **h** | 3-5 | **PCC (nuclear): effect of condition** | **one-way ANOVA** | <0.0001 | **** |

| | | | | | |
|---|---|---|---|---|---|
| | | Follow-up: mito-APEX2 vs. Su9-LbCas12a | Dunnett's multiple comparison test (corrected) | 0.8587 | ns |
| | | Follow-up: mito-APEX2 vs. COX8-LbCas12a | Dunnett's multiple comparison test (corrected) | 0.9930 | ns |
| | | Follow-up: mito-APEX2 vs. ATG4D-LbCas12a | Dunnett's multiple comparison test (corrected) | 0.9998 | ns |
| | | Follow-up: mito-APEX2 vs. None-LbCas12a | Dunnett's multiple comparison test (corrected) | 0.8825 | ns |
| | | Follow-up: mito-APEX2 vs. NLS-LbCas12a | Dunnett's multiple comparison test (corrected) | <0.0001 | **** |

| Figure 4 | | | | | |
|---|---|---|---|---|---|
| **Panel** | **Biological replicates** | **Comparison** | **Test** | ***P* value** | ***P* value summary** |
| **b** | 1-6 | **PCC (mito): effect of localization signal** | **Linear mixed effects model** | <0.0001 | **** |
| | | Follow-up: ATG4D vs. COX8 | Difference of least means squares (corrected) | <0.0001 | **** |
| | | Follow-up: ATG4D vs. mito-APEX2 localization signal | Difference of least means squares (corrected) | 0.313215 | ns |
| | | Follow-up: ATG4D vs. NLS | Difference of least means squares (corrected) | <0.0001 | **** |

| | | | | |
|---|---|---|---|---|
| | Follow-up: ATG4D vs. None | Difference of least means squares (corrected) | 0.092559 | ns |
| | Follow-up: ATG4D vs. Su9 | Difference of least means squares (corrected) | <0.0001 | **** |
| | Follow-up: COX8 vs. mito-APEX2 localization signal | Difference of least means squares (corrected) | 0.947609 | ns |
| | Follow-up: COX8 vs. NLS | Difference of least means squares (corrected) | <0.0001 | **** |
| | Follow-up: COX8 vs. None | Difference of least means squares (corrected) | <0.0001 | **** |
| | Follow-up: COX8 vs. Su9 | Difference of least means squares (corrected) | 0.088139 | ns |
| | Follow-up: mito-APEX2 localization signal vs. NLS | Difference of least means squares (corrected) | 0.009291 | ** |
| | Follow-up: mito-APEX2 localization signal vs. None | Difference of least means squares (corrected) | 0.197036 | ns |
| | Follow-up: mito-APEX2 localization signal vs. Su9 | Difference of least means squares (corrected) | 0.859550 | ns |
| | Follow-up: NLS vs. None | Difference of least means squares (corrected) | <0.0001 | **** |
| | Follow-up: NLS vs. Su9 | Difference of least means | <0.0001 | **** |

| | | | | |
|---|---|---|---|---|
| | | squares (corrected) | | |
| | Follow up: None vs. Su9 | Difference of least means squares (corrected) | <0.0001 | **** |

**Supplementary Table 2, Construction of Plasmids/Materials used**

| **Plasmid name** | **Protein Cassette** | **Cloning method** | **Sources** | **Notes** |
|---|---|---|---|---|
| pSKL.038 | None-Eco2 RT | PCR amplification | Eco2 RT from Addgene plasmid #184992[1] | |
| pSKL.039 | COX8-Eco2 RT | PCR amplification | | |
| pSKL.041 | None-Eco1 RT | PCR amplification | Eco1 RT from Addgene plasmid #184994[1] | |
| pSKL.043 | COX8-CasPhi | PCR amplification | CasPhi from Addgene plasmid #158801[2] | |
| pSKL.044 | COX8-LbCas12a | PCR amplification | | |
| pSKL.045 | COX8-CspRecT | PCR amplification | CspRecT from Addgene plasmid #138474[3] | |
| pSKL.046 | COX8-EcSSB | PCR amplification | EcSSB from Addgene plasmid #138474[3] | |
| pSKL.047 | COX8-BetRecT | PCR amplification | BetRecT from pKD46[4] | |
| pSKL.049 | COX8-Eco1 RT | PCR amplification | | |
| pSKL.050 | COX8-Cas3 | PCR amplification | Cas3 on gBlock, gift from Bondy-Denomy lab[5] | |
| pSKL.051 | COX8-Cas5 | PCR amplification | Cas5 on gBlock, gift from Bondy-Denomy lab[5] | |
| pSKL.052 | COX8-Cas7 | PCR amplification | Cas7 on gBlock, gift from Bondy-Denomy lab[5] | |

| | | | | |
|---|---|---|---|---|
| pSKL.053 | COX8-Cas8 | PCR amplification | Cas8 on gBlock, gift from Bondy-Denomy lab[5] | Entire line filtered out due to low cassette expression |
| pSKL.062 | COX8-PapRecT | PCR amplification | PapRecT from Addgene plasmid #138475[3] | |
| pSKL.063 | Su9-PapRecT | PCR amplification | | |
| pSKL.064 | ATG4D-PapRecT | PCR amplification | | |
| pSKL.065 | Su9-BetRecT | PCR amplification | | |
| pSKL.066 | ATG4D-BetRecT | PCR amplification | | |
| pSKL.067 | Su9-CspRecT | PCR amplification | | |
| pSKL.068 | ATG4D-CspRecT | PCR amplification | | |
| pSKL.069 | Su9-LbCas12a | PCR amplification | | |
| pSKL.070 | ATG4D-LbCas12a | PCR amplification | | |
| pSKL.071 | Su9-Cas3 | PCR amplification | | |
| pSKL.072 | ATG4D-Cas3 | PCR amplification | | |
| pSKL.074 | ATG4D-Cas8 | PCR amplification | | |
| pSKL.075 | COX8-SpyCas9 | PCR amplification | SpyCas9 from Addgene plasmid #184995[1] | |
| pSKL.076 | Su9-SpyCas9 | PCR amplification | | |
| pSKL.077 | ATG4D-SpyCas9 | Twist synthesis | | |
| pSKL.110 | NLS-CasPhi | PCR amplification | | |
| pSKL.111 | None-CasPhi | PCR amplification | | |

| | | | |
|---|---|---|---|
| pSKL.112 | NLS-LbCas12a | PCR amplification | NLS-LbCas12a from Addgene plasmid #126638[6] |
| pSKL.113 | None-LbCas12a | PCR amplification | |
| pSKL.114 | NLS-SpyCas9 | PCR amplification | |
| pSKL.115 | None-SpyCas9 | Twist synthesis | |
| pSKL.116 | None-BetRecT | PCR amplification | |
| pSKL.117 | None-CspRecT | PCR amplification | |
| pSKL.118 | None-PapRecT | PCR amplification | |
| pSKL.119 | None-Cas3 | Twist synthesis | |
| pSKL.127 | mito-APEX2 | Twist synthesis | mito-APEX2 from Addgene plasmid #72480[7] |
| pSKL.128 | Su9-EcSSB | Twist synthesis | |
| pSKL.129 | ATG4D-EcSSB | Twist synthesis | |
| pSKL.130 | Su9-Cas5 | Twist synthesis | |
| pSKL.131 | ATG4D-Cas5 | Twist synthesis | |
| pSKL.132 | Su9-Cas7 | Twist synthesis | |
| pSKL.133 | ATG4D-Cas7 | Twist synthesis | |
| pSKL.134 | Su9-Eco1 RT | Twist synthesis | |
| pSKL.135 | ATG4D-Eco1 RT | Twist synthesis | |
| pSKL.136 | Su9-Eco2 RT | Twist synthesis | |
| pSKL.137 | ATG4D-Eco2 RT | Twist synthesis | |
| pSKL.138 | COX8-SaCas9 | Twist synthesis | |
| pSKL.139 | Su9-SaCas9 | Twist synthesis | |
| pSKL.140 | ATG4D-SaCas9 | Twist synthesis | |
| pSKL.141 | NLS-SaCas9 | Twist synthesis | |
| pSKL.142 | COX8-PlmCasX | Twist synthesis | PlmCasX from Addgene plasmid #180512[8] |
| pSKL.143 | Su9-PlmCasX | Twist synthesis | |

| | | | |
|---|---|---|---|
| pSKL.144 | ATG4D-PlmCasx | Twist synthesis | |
| pSKL.145 | NLS-PlmCasX | Twist synthesis | |
| pSKL.146 | COX8-DpbCasX | Twist synthesis | DpbCasX from ref.[8] |
| pSKL.147 | Su9-DpbCasX | Twist synthesis | |
| pSKL.148 | ATG4D-DpbCasX | Twist synthesis | |
| pSKL.149 | NLS-DpbCasX | Twist synthesis | |
| pSKL.150 | COX8-PaSSB | Twist synthesis | |
| pSKL.151 | Su9-PaSSB | Twist synthesis | |
| pSKL.152 | ATG4D-PaSSB | Twist synthesis | |
| pSKL.153 | COX8-MsSSB | Twist synthesis | |
| pSKL.154 | Su9-MsSSB | Twist synthesis | |
| pSKL.155 | ATG4D-MsSSB | Twist synthesis | |
